# The Structure of Escherichia coli MscL and its dimer formation in Nanodiscs

**DOI:** 10.64898/2026.07.06.736777

**Authors:** Tim Rasmussen, Julia Isabel Bahner, Vanessa J. Flegler, Tamsanqa T Hove, Christian Kraft, Akiko Rasmussen, Bettina Böttcher

## Abstract

Mechanosensitive channels of large conductance (MscL) are essential bacterial safety valves that prevent osmotic lysis by releasing solutes in response to membrane tension. Despite extensive functional studies on *Escherichia coli* MscL (EcMscL), its high-resolution structure remained unknown. Using cryo-electron microscopy, we present an experimental structure of EcMscL reconstituted in nanodiscs at 3.1 Å resolution. The structure reveals a pentameric assembly with a narrow hydrophobic gate at the cytosolic side and a periplasmic cavity, consistent with the canonical MscL-fold. Differences to earlier published crystal structures of MscL from other organisms are in the less conserved periplasmic loop. We observe a previously unreported dimeric association of EcMscL pentamers, mediated by residues 61-63 in the periplasmic loop. This dimeric interface is located at the periplasmic side and provides a structural basis for the formation of higher-order clusters. The observed arrangement enables a fluid-like, mosaic packing of channels with center-to-center distances of 5.9–9 nm, consistent with biophysical and imaging data. These findings provide a structural framework for understanding cluster organization of EcMscL that modulates its activity in cellular stress response.

## Introduction

Bacteria experience many life-threatening, environmental challenges. One of them is a hypoosmotic shock with a sudden drop in osmolytes. Such an event leads to a rapid influx of water followed by an increase in the turgor pressure and eventually membrane rupture in the absence of protective mechanisms. Bacteria are protected against membrane rupture by mechanosensitive channels, which gate when the membrane tension exceeds certain thresholds. The open channels allow solutes to equilibrate between inside and outside of the cell and release the osmotic pressure.

One family of protective channels are mechanosensitive channels of large conductance (MscL), which have conductivities in the range of 3-4 nS [1-3]. These channels gate at membrane tensions close to the point of rupture with an estimated pore diameter of 2-4 nm [4-6]. Thus, opening of MscL leads to a rapid loss of many solutes including small proteins. As such, the protection by MscL channels is costly for the organism and can be regarded as a last resort to allow for the cell survival.

Structural data of MscL has been obtained from few crystal structures of various species [7-12]. The best resolved structure so far represents MscL from *Mycobacterium tuberculosis* (MtMscL) [7, 8]. MtMscL has a pentameric arrangement of subunits in which each subunit consists of a short N-terminal amphiphilic helix, two transmembrane helices (TM1, TM2) and a C-terminal helix. Together, the TM1s and TM2s generate a water filled cavity on the periplasmic side of the channel. On the cytosolic side the TM1s provide a narrow hydrophobic pore. The C-terminal helices assemble into a helical bundle that is tethered via linkers to the cytosolic side of the channel. This C-terminal helical bundle is dispensable [13, 14] for gating but together with the linkers might provide a molecular sieve that restricts access to the pore [14, 15].

Structural insights into how MscL opens comes from other MscL-like channels of the archaeon *Methanosarcina acetivorans* (MaMscL) [10] and from the gram-positive bacterium *Staphylococcus aureus* (SaMscL) [9]. Both structures show an expanded state with a wider pore and tilted TM-helices that generate a shortened membrane spanning width. Despite the different oligomeric states of tetrameric SaMscL and pentameric MaMscL, the narrowest pore diameter is almost identical and with some 0.8 nm [10] much tighter than the expected open pore diameter of 2-4 nm based on release assays [4, 5]. Therefore, it is likely that these crystal structures represent intermediate states, which then progress into the fully open state by outward sliding of the subunits following the two-step helix-pivoting model [9]. This model is supported by many all-atom molecular dynamics (MD) trajectories and finite element calculations that are either steered by pulling on the N-terminal helix or by changing the membrane tension [6, 16-20].

To understand gating of MscL and its function it is necessary to consider its action in higher order aggregates, which have been observed by fluorescence microscopy, small-angle neutron scattering, atomic force microscopy and neutron reflection [21-23]. In these clusters, channels act cooperatively and their response depends on the proximity to other channels [21]. Theoretic considerations based on continuum mechanics support these experimental findings [24]. Today, most functional data were obtained from MscL of *Escherichia coli* (EcMscL) for which no experimental structure has been available. All these studies assume structural similarity to MtMscL. To confirm that this assumption is correct and to allow a more detailed understanding on side chain level we have used electron cryo microscopy (cryo-EM) and single particle image processing to determine an experimental structure of EcMscL. We found that the channels associate further into dimers of pentamers suggesting a structural route for cluster formation in membranes.

## Results

We purified EcMscL via a C-terminal His-tag followed by size exclusion chromatography (SEC) (Figure 1, Figure S1). EcMscL eluted in two main peaks at 9.7 ml and 11.6 ml, which correspond to molecular masses of 370 kDa (peak 1) and 160 kDa (peak 2) according to the SEC mass calibration. We reconstituted EcMscL of both peaks into nanodiscs for further structural studies by electron cryo-microscopy (cryo-EM). For both reconstitutions, cryo-EM and 2D-classification showed evenly distributed elongated nanodiscs of similar sizes with one or two channels per disc (Figure 1). One MscL per nanodisc was the predominant species in both reconstitutions, with two MscL per nanodisc being more frequent in the reconstitution of peak1.

**Figure 1.**
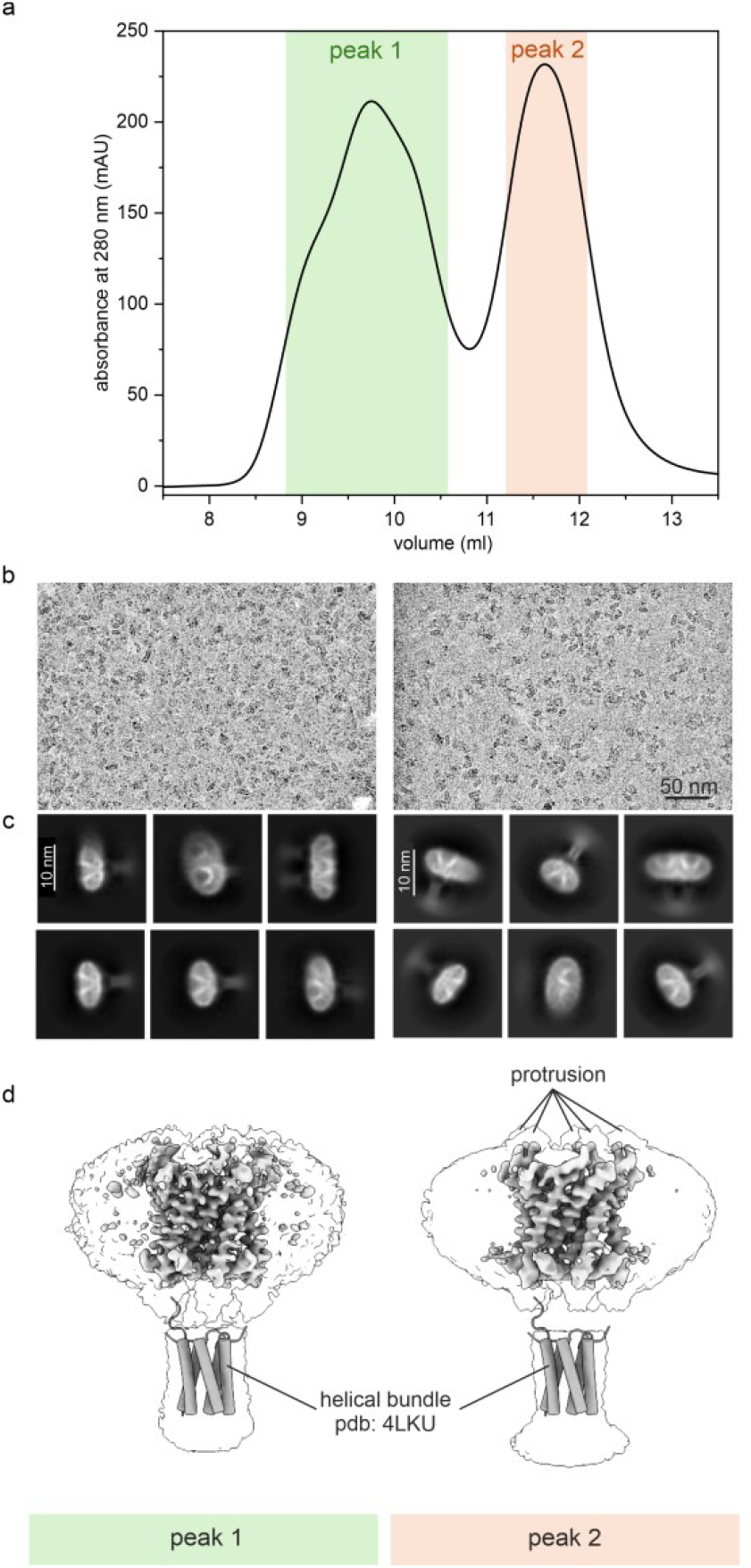
Characterization of EcMscL. a) Size exclusion chromatography of solubilized EcMscL. EcMscL elutes in two peaks from a Superdex 200 Increase 10/300 GL column at 9.7 ml (peak 1) and at 11.6 ml (peak 2). **b)** Part of micrographs of vitrified EcMscL in nanodiscs showing an even distribution of particles. **c)** Selected class averages of nanodiscs with one or two channels. **d)** Surface representations of the 3D-maps at a low threshold (translucent) and at a high threshold (solid). The crystal structure of the C-terminal domain of EcMscL (4LKU [11]) was fitted and is shown as tubes.

Ab-initio reconstructions with C1-symmetry revealed a pentameric MscL positioned out of centre close to the edge of the of the nanodiscs. This position of MscL in the nanodisc was well defined as the surrounding helical membrane scaffolding protein was clearly resolved in contrast to the opposite side, which was more variable (Figure S2). The initial maps were further refined with imposed C5-symmetry to resolutions of 3.1 Å (peak 1) and 3.5 Å (peak 2), respectively. Both maps resolved the N-terminal membrane-embedded part to high resolution with clear side chain density in most regions. In contrast, the C-terminal helical bundle was only visible at a lower threshold without resolving the individual helices (Figure 1). This suggested mobility or variability of the bundle in respect to the membrane part.

We modelled most of the membrane embedded part of MscL automatically with ModelAngelo [25], which further highlighted the quality and interpretability of the maps in the transmembrane region. The refined models of the reconstitutions of peak 1 and of peak 2 comprised residues 2-104 and were identical (RMSD=0.9 Å, Figure S3) within the expected errors of modelling [26]. Both models revealed a pentameric assembly of subunits with a narrow gate on the cytosolic side and an accessible cavity on the periplasmic side (Figure 2).

**Figure 2.**
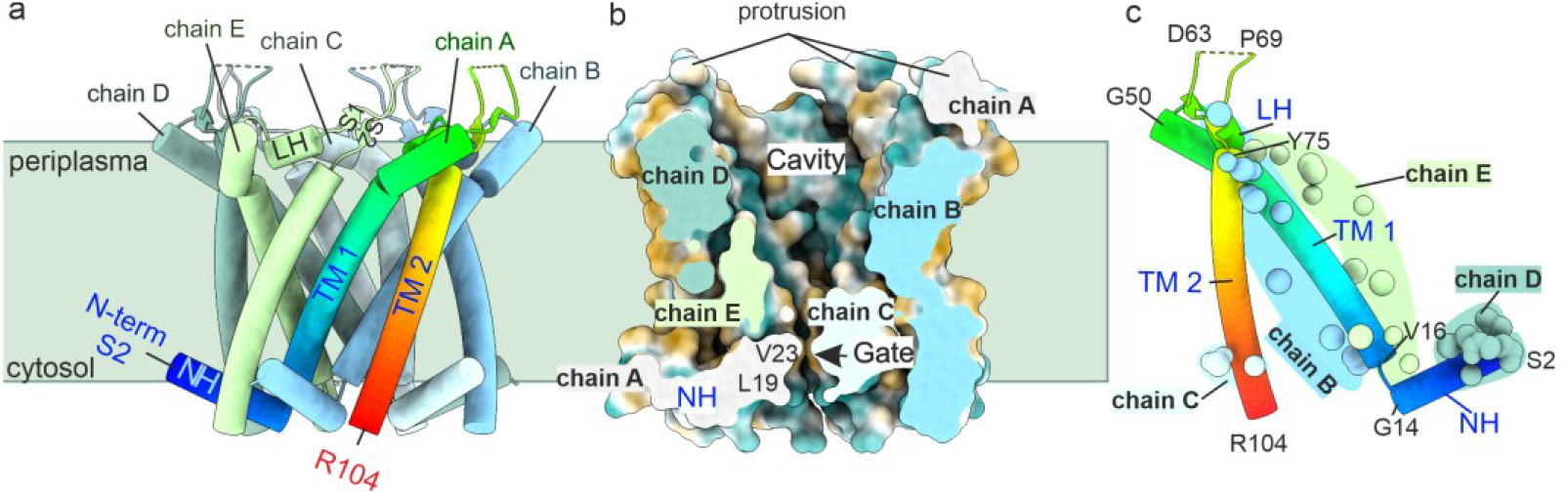
Model of EcMscL and contact to other subunits. **a)** EcMscL in cartoon representation with helices shown as tubes. One of the subunits is coloured in rainbow from blue at the N-terminus to red at the last resolved residue at the C-terminus (R104). The other subunits are shown in different colours. Both, N- and C-termini are located on the cytosolic side. The approximate position of the membrane is indicated by a green square. **b)** The model of EcMscL is presented as Van-der Waals surface and coloured according to the hydrophobicity (from brown for hydrophobic to blue for hydrophilic). The front half is clipped off to reveal a cross-section along the channel axis. The clipping plane is coloured according to the identity of the respective subunit and shows intersection with all subunits. At the periplasmic side the channel forms a water accessible cavity and on the cytosolic side a narrow, hydrophobic gate. Periplasmic loops between TM1 and TM2 protrude from the plane of membrane. **c)** A single subunit of MscL is coloured in rainbow. The spheres highlight atoms in adjacent subunits within 3.5 Å. The subunit interacts with all other subunits in the channel. Residues 64-68 in the periplasmic loop are unresolved and mark the tips of the protrusions.

The EcMscL-subunits consisted of an N-terminal amphiphilic helix (NH, residues 1-14) and two transmembrane helices (TM1, residues 16-50 and TM2, residues 75-104) connected by a periplasmic loop (residues 51-74) (Figure 2). We could not model the C-terminal residues 105-136 that form a linker and the C-terminal helix. In addition, residues 64-68 of the periplasmic loop were unresolved but located to the protrusions on the periplasmic surface. The fold of EcMscL is the same as the one of MtMscL determined by X-ray crystallography (Figure S7) [7] .

The fold of the EcMscL subunit was extended so that each subunit interacted with all other subunits within the channel (Figure 2) but had only few interactions within a subunit. The intra-subunit interactions were provided mainly by hydrophobic contacts interconnecting TM1 and TM2 close to the centre of the membrane (Figure 3a). Other intra subunit contacts located to the periplasmic loop, where a short helix (LH, D53-F57) provided a central hub that formed hydrophobic interactions with TM1 and with TM2 (Figure 3a).

**Figure 3.**
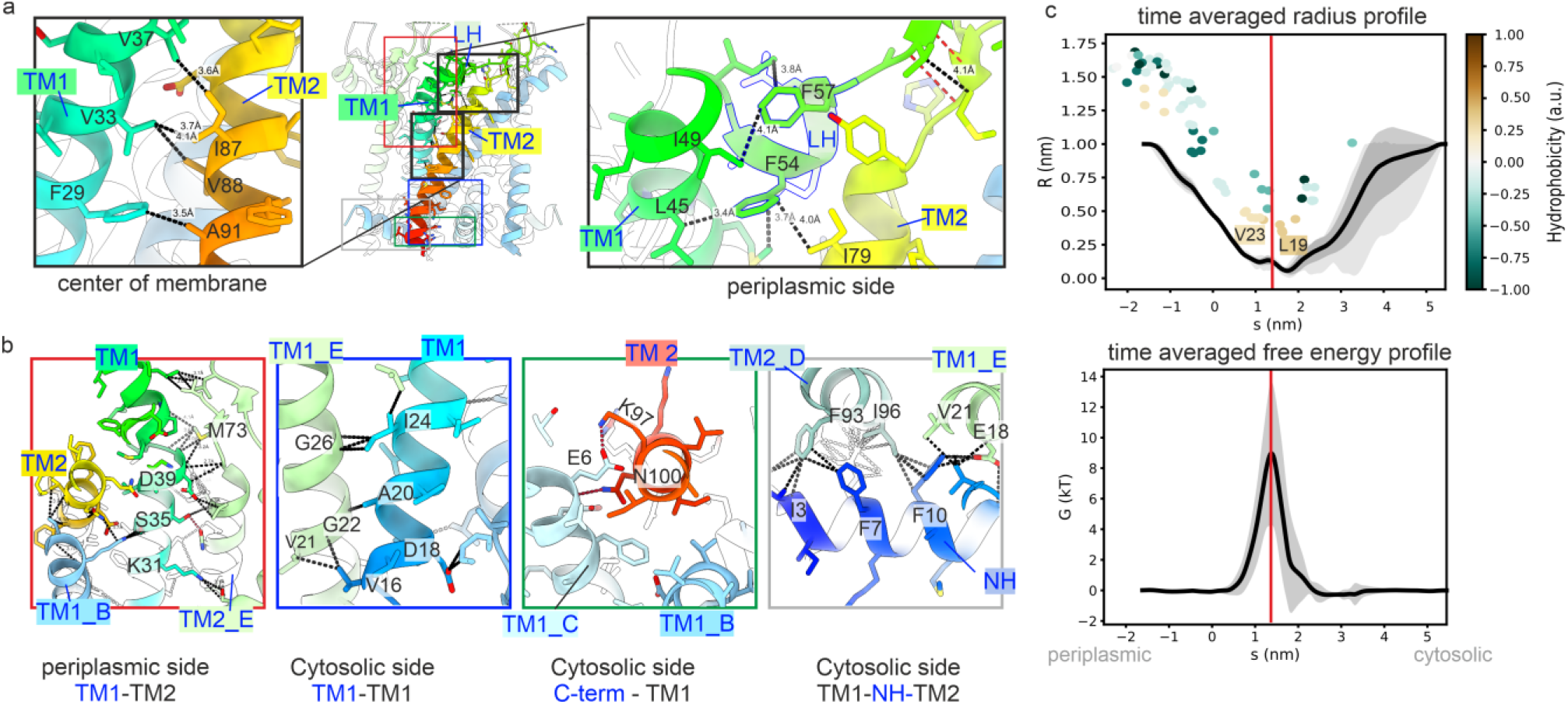
Intra and inter-subunit interactions: **a)** Specific intra-subunit interactions. The left panel shows the hydrophobic interactions between TM1 and TM2 and the right panel the interactions of the loop helix (LH) in the periplasmic loop with TM1 and TM2. The central panel gives an overview of the whole channel with the same colouring of the subunits as in the close-ups. The approximate positions of the close-ups are indicated by black squares for the intra-subunit interactions in **a)** and with coloured squares for the inter-subunit interactions shown in **b). b)** Inter-subunit interactions with chain A as reference chain. Contacts are shown as dotted lines. The residue numbers of interacting residues are labelled. **c)** Channel annotation of short part of a MD-trajectory (21-23 ns) with CHAP [27]: The solvent density in the penetration path for the full trajectory is shown in Figure S5a. The top panel shows the time averaged radius profile. S specifies the position along the channel axis and R the pore-radius. The probe for the pore radius estimation is truncated at 1 nm. Channel lining residues are shown as spheres and coloured according to their hydrophobicity. The colour key shows the hydrophobicity in arbitrary units from brown=hydrophobic to blue hydrophilic. The residues V23 and L19 define the narrowest part of the pore. The red line denotes the position of the maximal energetic barrier for water molecules to pass. The lower panel shows the free energy for water molecules to penetrate the channel. The largest energetic barrier for water is between residues L19 and V23 (red line). This position does not change in later frames of the trajectories (see Figure S5 for energy plot at 101-103 ns).

Cross-subunit interactions were more frequent (Figure 3b). On the periplasmic side, TM2 together with the periplasmic loop made cross-subunit contacts with TM1. On the cytosolic side, the TM1s were tightly packed with numerous interhelical contacts close to the narrowest constriction of the pore. Other inter-subunit interaction hot spots were the C-terminus of TM2 and the N-terminal helix. Here, TM2 formed hydrophobic interactions, H-bonds and salt bridges with the N-terminal helices on both sides. A salt bridge was evident between K97 with E6 and an H-bond between N100 and the backbone of the N-terminal helix.

To further analyse the model, we calculated three short MD trajectories with a length of 130 ns, 200 ns and 200 ns respectively (n=3). EcMscL was remarkably stable and showed little fluctuations in the backbone positions, which agreed with the well resolved details in the EM-map of the membrane embedded part (Figure S4). In contrast, the tips of the periplasmic loop at the protrusions were highly mobile, which agreed with the failure to resolve residues 64-68 in the EM-maps. H-bonds between the C-terminus of TM2 involving N100 and the N-terminal helix on the adjacent subunits involving E6 and K97 were fluctuating but were maintained throughout the simulation. In addition, K97 also coordinated the phosphate-group of an adjacent lipid (Figure S4). After formation of this interaction, it remained stable to the end of the simulation as shown for one of the MD-trajectories Figure S4). Although K97 coordinated a lipid headgroup, lipids did not enter the inter-subunit interface between the N-terminal helix and TM2 on this side. On the other side of the N-terminal helix, lipids entered more readily the inter-subunit space to TM2 (Figure S4). In general, lipids maintained positional mobility without binding to confined pockets. This agreed with the EM-maps that did not resolve lipid like densities associated with MscL.

We further analysed short sub-trajectories of 2 ns (21-23 ns, Figure 3; 101-103 ns Figure S5) to identify the penetration pathway with the program CHAP [27, 28]. Residues L19 and V23 formed two hydrophobic rings at the cytosolic side of the pore and provided the narrowest constriction in the penetration pathway with a diameter of 0.10±0.03 nm (Figure 3c). This gate was impermeable for water with the maximum of the calculated energetic barrier being between the two hydrophobic rings. All three MD-trajectories showed that the hydrophobic lock was maintained throughout the simulation as by the low water density at the gate (Figure S5).

We were intrigued by the fact that many nanodiscs contained two channels per disc. As both channels were visible in class averages, we reasoned that their association was not random and we attempted to obtain a three-dimensional map of the assembly. Ab-initio models without imposed symmetry showed one of the two channels completely and the second channel partly (Figure S6). This was sufficient to unambiguously fit the two channels as rigid bodies. Further processing with C2-symmetry resolved both channels at intermediate resolution identifying the helical packing (Figure 4). The channels were oriented such that at the periplasmic side two TM1s faced each other and that the periplasmic loops from the adjacent subunits came into close contact at residues 61-63. The dimeric contact involved four subunits (Figure 4a). Rigid body placement of the EcMscL model suggested that salt bridges between R62 and D63 could stabilize the cross-channel interaction via the periplasmic loops after subtle movements of the loops. However, in the model generated by rigid body placement, the respective side chains were some 7 Å apart, which was too far for the formation of a salt bridge. In addition, hydrophobic contacts between L61 with L48 in TM1 on another subunit contributed a hydrophobic cross-channel interaction. In contrast to the periplasmic side, the channels did not have direct protein-protein contacts on the cytosolic side.

**Figure 4.**
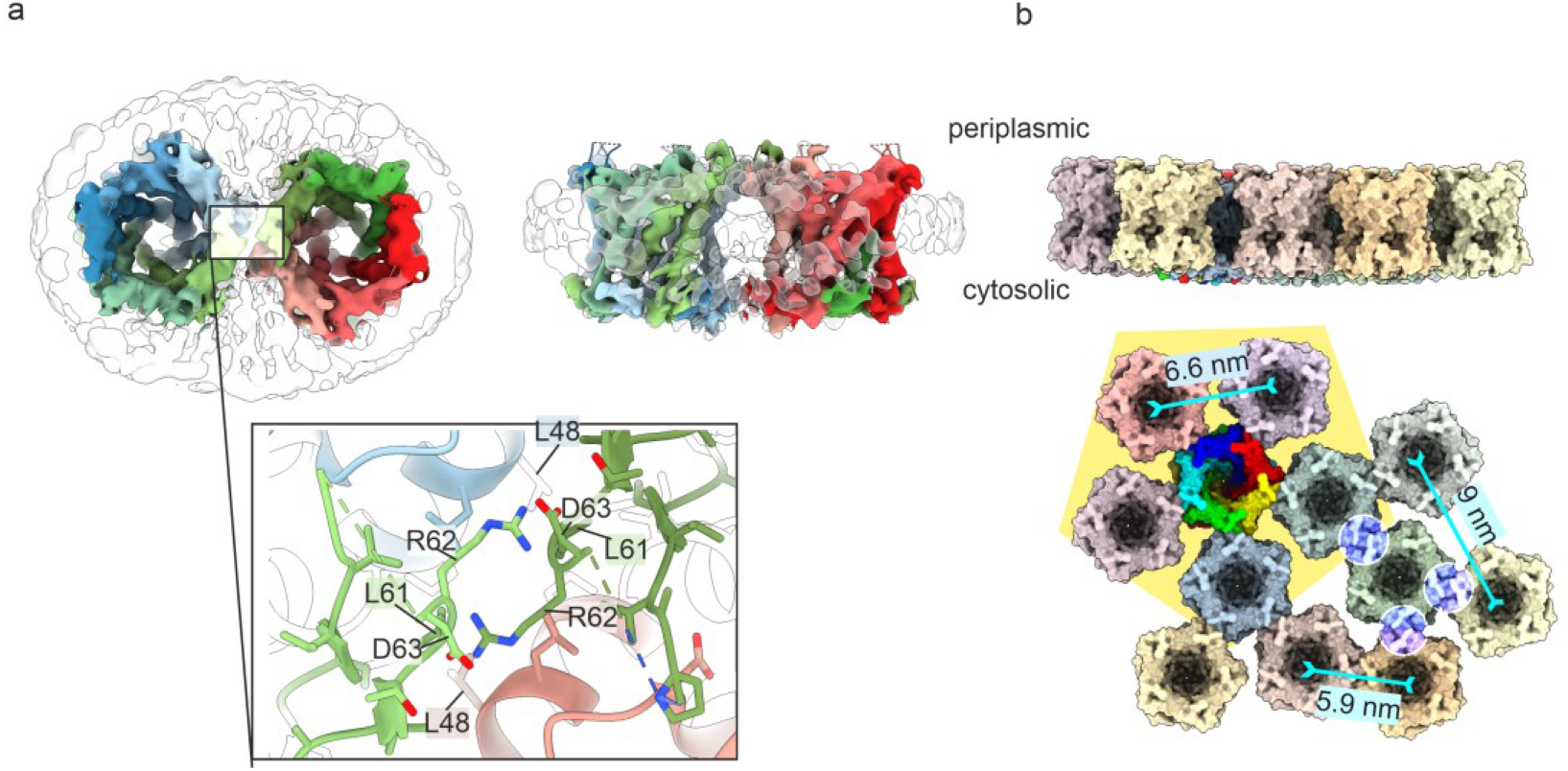
Dimeric association of EcMscL. **a)** Map and model of two EcMscL channels in a nanodiscs. The EcMscL dimers are viewed along the dimer axis from the periplasmic side (left) and perpendicular to the dimer-axis (right). The surface of the map is shown at a lower threshold as translucent outline and at a higher threshold coloured within 3.5 Å of the placed models. The close-up shows the interaction side with the subunits coloured differently. L48 in TM1 and L61 in the periplasmic loop of the opposite MscL form hydrophobic contacts. R62 and D63 in the opposing periplasmic loops are positioned favourably to form a salt bridge but would require slight adjustments of the loop to bring them into direct contact. **b)** Cluster of EcMscL channels generated by maintaining the dimer interactions. In one of the channels the subunits are coloured in rainbow. This channel maintains dimer contacts with all five surrounding channels without clashes (subcluster with yellow background). Channels in the periphery of this sub-cluster can only maintain up to three dimeric interactions without clashes. These three dimer contacts are encircled in one of the channels. The centre-to-centre distance of channels with a dimeric contact is 5.9 nm. Nearest neighbours without dimeric contact have centre-to-centre distances of 6.6 nm and 9 nm.

To test whether the arrangement of dimers of channels would remain stable in a membrane we obtained three short MD-trajectories (201 ns, 171 ns and 102 ns) of the rigid body dimer model placed into a membrane. In all three trajectories, the channels remained closely associated throughout the simulation time. Next, we asked, whether H-bonds formed between R62 and D63 of one channel with the same residues in the opposite channel. In all three trajectories such an H-bond formed quickly. However, over time the H-bonds fluctuated and only rarely both residues, R62 and D63, formed bonds with the opposing channel at the same time (Figure S6). These simulations suggested that the small contact side was sufficient to keep two channels associated in a dimer. To test *in silico* whether the dimeric arrangement could support the formation of larger clusters, we placed several EcMscL in proximity to generate as much dimer contacts as possible without clashing (Figure 4b). In general, pentamers cannot form extended, planar arrays with equal contacts to other pentamers and are typically associated with curved assemblies. However, a central pentamer supported dimeric contacts with five surrounding pentamers without clashing (Figure 4). The centre-to-centre distances of the outer pentamers to the central pentamer was 5.9 nm. The pentamers in the outer periphery did not touch their neighbours and had a centre-to-centre distance of 6.6 nm. These outer pentamers supported two more dimeric contacts without clashing giving a total of three possible dimeric contacts per pentamer in this shell. There are several possibilities which subunits within a pentamer form a dimeric contact. Therefore, the packing of channels in a cluster was a variable, fluid-like mosaic with typical centre-to-centre distances between the nearest neighbours of 5.9 nm in a dimer, and 6.6 nm and 9 nm in neighbours without direct protein-protein contact. Such a cluster is almost planar.

## Discussion

Most functional information of MscL channels has been obtained from EcMscL and numerous MD-simulations have modelled possible structural responses of EcMscL based on homology models. Here we provide an experimental structure of EcMscL in nanodiscs. The structure closely resembles that of *M. tuberculosis* MscL (MtMscL), consistent with their high sequence similarity. Indeed, fitting the crystal structure of MtMscL [7], as a rigid body into our EM-map shows good agreement but also reveals marked differences at the periplasmic side (Figure S7): Compared to MtMscL, the C-terminal end of TM1 of EcMscL is bent outward and the tip of the periplasmic loop protrudes out of the plane of membrane. In contrast, TM1 of MtMscL reaches the periplasmic side closer to the symmetry axis (smaller radii) and the periplasmic loop of MtMscL forms extensive interactions with the neighbouring subunit. Consequently, the cross-section of MtMscL on the periplasmic side is almost circular whereas it has a pronounced pentagonal shape in EcMscL.

The structural differences of MtMscL and EcMscL on the periplasmic side mirror the lower sequence conservation of the periplasmic loop and at the C-terminal end of TM1 compared to the rest of the protein (Figure S7).

It is conceivable that the observed dimerization in some nanodiscs arises from non-specific, concentration-driven aggregation during reconstitution. Such a scenario would likely favour random associations between channels, with a preference for edge-to-edge contacts to minimize the exposed perimeter and maximize packing efficiency within the nanodisc. However, we observe a specific, vertex-oriented dimer interface, which is mediated by defined interactions in the periplasmic loop, suggesting a more structured association, rather than a random association as consequence of high protein concentration. The periplasmic loop provides the cross-channel interaction between EcMscL channels. MD simulations suggest that although these interactions are variable (Figure S6), they are sufficient to keep the channels associated in a membrane. It is conceivable that such cross-channel interactions drive the association of EcMscL into larger clusters. Such clusters have been observed by atomic force microscopy (AFM), small angle neutron scattering and fluorescence microscopy [7]. Estimations of distances from AFM suggest a centre-to centre distance of 5 nm, which is similar to the distance of 6 nm that we observe in the dimers. Although, the interaction interface in the dimers is small, it would be sufficient to steer a preferential way of channel association across the vertices (Figure 4b).

Such association generates membrane filled cavities between the channels and prevents a dense hexagonal packing as observed in 2D-crystals [29]. The cavities provide space for channels to expand during opening. Calculations of elastic potential suggest that clustering reduces the tension required for opening and increases the open probability [24]. The estimated centre-centre distances to the nearest neighbours in a cluster of 6-9 nm (Figure 4) is within the range where a cooperative effect for gating is expected based on the elastic potential calculations [24].

In the clusters, the dimeric contacts generate fix points on the periplasmic side. Further radial expansion across these fix-points would require a cooperative increase in the centre-to-centre distances for opening of a neighbouring channel. A recent structural model of an expanded EcMscL in DSPC nanodiscs shows a marked increase in the diameter across the TM1-vertices on the periplasmic side [30]. In a cluster, such radial expansion of the vertices would affect the cluster contacts and impact their chemical properties. Indeed, ssNMR of a mutant (G22S), shows chemical shift differences at the contact residues R62 and D63 in liposomes while surrounding residues where not resolved in the mutant [31].This observation makes an unexpected connection between the dimer contact and a distant mutation close to the gate that leads to gating at lower tension similar as expected for EcMscL in a cluster. It also suggests that the contact is maintained in the mutant while the surrounding loop is more flexible.

Within a cluster it appears unlikely that several adjacent channels can expand across the vertices. However, the loose fluid like packing of the channels with unsatisfied vertex contacts provides space for a subset of distant channels to expand without the necessity of larger reorganization of the whole cluster. Consequently, the cooperative coupling between the channels is disturbed by the irregular arrangement and only a subset of channels is likely to open. This fits with the observation that only a small percentage of channels in *E*.*coli* appears active in gating at a given time [23].

In conclusion, the dimeric association of EcMscL provides a structural framework to understand how channels form larger clusters that maintain close association required for cooperative gating and provide space between channels to enable area expansion of a subset of channels.

## Materials and methods

### Expression, purification and reconstitution into nanodiscs of EcMscL

The plasmid pET-MscL-His_6_ was transformed into the *Escherichia coli* strain BL21(DE3). An overnight culture in LB medium was used to inoculate a pre-culture and then a main culture in LB medium at 37 °C. At an OD_600_ of 1 expression of MscL was induced with 1 mM IPTG and continued to grow for 4 hours at 37 °C. The cells were harvested by centrifugation at 5000 g for 30 min and stored at –80 °C until further use. The cells were solubilised for 1 h on ice with a buffer containing 50 mM sodium phosphate buffer pH 7.5, 300 mM NaCl, 10 % (w/v) glycerol, 25 mM imidazole, 0.2 mM PMSF, 5 mM EDTA, 1.6 mg/ml lysozyme, and 1.5 % DDM. After centrifugation at 7200 g at 4 °C for 45 min, the supernatant was applied onto a gravity-flow column prepared with a Ni-NTA bed and washed (50 mM sodium phosphate buffer pH 7.5, 300 mM NaCl, 25 mM imidazole, 0.5 % DM). Subsequently, His-tagged MscL was eluted using buffer containing 50 mM sodium phosphate buffer pH 7.5, 300 mM NaCl, 300 mM imidazole, and 0.5 % DM. The MscL-containing elution fraction was further purified via size exclusion chromatography using a Superdex 200 Increase 10/300 GL column (Cytiva) in buffer containing 150 mM HEPES pH 7.5, 50 mM NaCl, 5 mM EDTA and 0.15 % DM. Reconstitution into MSP-1E3D1 nanodiscs was done as described previously for MscS [32]. Briefly, DDM-purified MscL, Azolectin (in 20 mM Tris pH 7.5, 100 mM NaCl, 0.5 mM EDTA, 200 mM sodium cholate), and MSP-1E3D1 (in 20 mM HEPES pH 7.5, 100 mM NaCl, 0.5 mM EDTA, 50 mM sodium cholate) were mixed at a 1:8:80 molar ratio and incubated for 2 h at room temperature. Biobeads were added and the sample was incubated overnight at 4 °C. Next day, another size exclusion chromatography was performed using the same buffer as previously with the detergent omitted. Purifications were monitored via SDS-PAGE using self-made 15 % gels. For Western blot detection, a Penta-His HRP conjugate antibody (Qiagen) against the C-terminal His6 tag of the construct was employed. Samples were concentrated to 1 mg/ml with Amicon 100 K Ultra-0.5 centrifugal filter device (Millipore) for cryo-EM sample preparation.

### Cryo-EM and structural data analysis

#### Vitrification

For cryo-EM, Quantifoil UltrAU Foil grids (300 mesh, R1.2/1.3) were glow-discharged in air for 2.5 min at medium power in a Harrick PDC-002 plasma cleaner. Samples were vitrified with a Vitrobot IV (FEI) with the humidity in the chamber set to 100 % and the temperature to 4 °C. 3.5 μl of the sample were applied onto the grids, excess liquid was blotted for 4 s with the blot force set to -20. Subsequently, the grids were plunge-frozen in liquid ethane cooled to liquid nitrogen temperature. Vitrified samples were stored in liquid nitrogen until further usage.

#### Data acquisition

Vitrified grids of samples were transferred to a Titan Krios G3 equipped with a Falcon IVi camera and a Selectris X energy filter. Zero-loss movies were collected with a slit width of 5 eV, at a nominal magnification of 130,000 (calibrated pixel size of 0.946 e^-^/Å^2^) with an exposure of 70 e^-^/Å^2^ in EER-format [33]. Movie acquisition was automated with EPU (Thermo Fisher) with the fast option using beam tilt corrected image shift. The conditions for Cryo-EM data collection are summarized in table S1.

#### Processing

Movies were dose-weighted and motion-corrected during the CryoSPARC [34] live session. Other preprocessing steps were patch CTF-estimation, blob-based particle picking, and extraction of particles in a 256 x 256 px^2^ box cropped to 128x128 px^2^. The cropped particles were subjected to streaming 2D-Classification. Particles from the best classes were selected to further rounds of 2D classification and selection. Image processing was continued within CryoSPARC following general procedures as summarized in the processing scheme (Figures S8 and S9). While ab-initio determination of 3D-maps gave reasonable results, these maps failed as references in iterative refinement. Therefore, we used the ab-initio approach for high resolution structure determination [35] by optimizing the resolution band between 3 Å and 6 Å, using a small Fourier radius step of 0.004 and larger initial and final minibatch sizes of 300 and 900. The best ab-inito 3D-classes were further refined by local refinement with a small rotation search extend of 2° and a small shift search of 1 Å. Angle search and shift search were recentered at each iteration. The initial low pass filter of the local search was 6 Å.

#### Model Building

Maps were interpreted and subjected to de novo Model building with Modelangelo [25]. ModelAngelo models of MscL in nanodiscs were manually corrected and completed in Coot [36] and further refined by real-space refinement in Phenix [37, 38]. [36-39]. The validation statistics are summarized in table S1.Figures of maps and models were generated with ChimeraX [40].

#### Channel annotation

For molecular dynamics (MD) simulations, the models were placed into a membrane bilayer of 60% POPE, 20% POPG and 20% DSPS in water with 150 mM KCl using CHARMM-GUI [41]. The CHARM-Gui-Input generator [42] generated the GROMACS [43] production run. The derived GROMACS inputs were used with the default parameters, apart from the initial minimization (changed: nsteps from 5000 to 100000; and emtol from 1000 to 500), because energy minimization was frequently not completed within the first 5000 steps. The barostat was C-rescale with semi-isotropic coupling and the thermostat was v-rescale at a temperature of 303.15 k. H-bonds were constraint with LINCS during all steps (minimization, equilibration and production). The time step was 1 fs during the first 0.375 ns of equilibration and 2 fs afterwards. Electrostatic interactions were calculated with PME and Verlet was used as cut-off scheme. The full parameter list of the production run is listed in the supplements. The MD-simulation was repeated 3 times (n=3) with production runs of 130 ns, 200 ns and 200 ns for the channel monomer and 102 ns, 171 ns and 201 ns for the dimer of channels.. The MD-trajectories were further analyzed with CHAP [27, 28] to identify pore properties such as hydrophobicity, pore diameter and water permeability.

## Data Availability

EM-maps were deposited in the Electron Microscopy Data Bank (EMDB). The corresponding model was deposited in the RCSB Protein Data Bank (RCSB PDB) with the accession codes: EMD-56958, PDB-28YC (EcMscL pentamer in nanodiscs); EMD-58096 (EcMscL dimer of pentamers in nanodiscs).

## Acknowledgements

Cryo-electron microscopy was carried out in the cryo-EM facility of the Julius-Maximilians-Universität Würzburg funded by the Deutsche Forschungsgemeinschaft (DFG, German Research Foundation – Projects INST 93/903-1 #359471283, INST 93/1042-1 #456578072, INST 93/1143-1 # 525040890). B.B. acknowledges project funding by the DFG (BO1150/15-2 # 343886090,BO1150/21-1 # 538122946 and BO1150/23-1 # 583559912).

## Author contributions

T.R., A.R. V.J.F. and B.B. developed the experimental design. V.J.F., T.R: J.I.B. purified EcMscL. C.K., T.R. and V.J.F. vitrified samples. C.K., B.B. and T.R. acquired electron microscopic data. B.B. and T.H. did electron microscopic image processing. T.R: and B.B. did model-building. B.B., T.R. and J.V.F. designed the figures. T.R., A.R., V.J.F. and B.B. interpreted the data. All authors contributed to writing and editing of the manuscript.

