## Supplemental Information for "The Structure of Escherichia coli MscL and its dimer formation in Nanodiscs"

### MD-simulation Production run- Parameters

The MD-simulation parameter files for Gromacs [1] were generated with CharmGui [2]. The parameters of the production runs are given below and was used for the monomers and dimers.

```
integrator            =md
dt                    =0.002
nsteps                =500000
nstxout-compressed    =50000
nstxout               =0
nstvout               =0
nstfout               =0
nstcalcenergy         =100
nstenergy             =1000
nstlog                =1000
cutoff-scheme         =Verlet
nstlist               =20
rlist                 =1.2
vdwtype               =Cut-off
vdw-modifier          =Force-switch
rvdw_switch           =1.0
rvdw                  =1.2
coulombtype           =PME
rcoulomb              =1.2
tcoupl                =v-rescale
tc_grps               =SOLUMEMBSOLV
tau_t                 =1.01.01.0
ref_t                 =303.15303.15303.15
pcoupl                =C-rescale
pcoupltype            =semiisotropic
tau_p                 =5.0
compressibility       =4.5e-54.5e-5
ref_p                 =1.01.0
constraints           =h-bonds
constraint_algorithm       =LINCS
continuation          =yes
nstcomm               =100
comm_mode             =linear
comm_grps             =SOLU_MEMBSOL
```

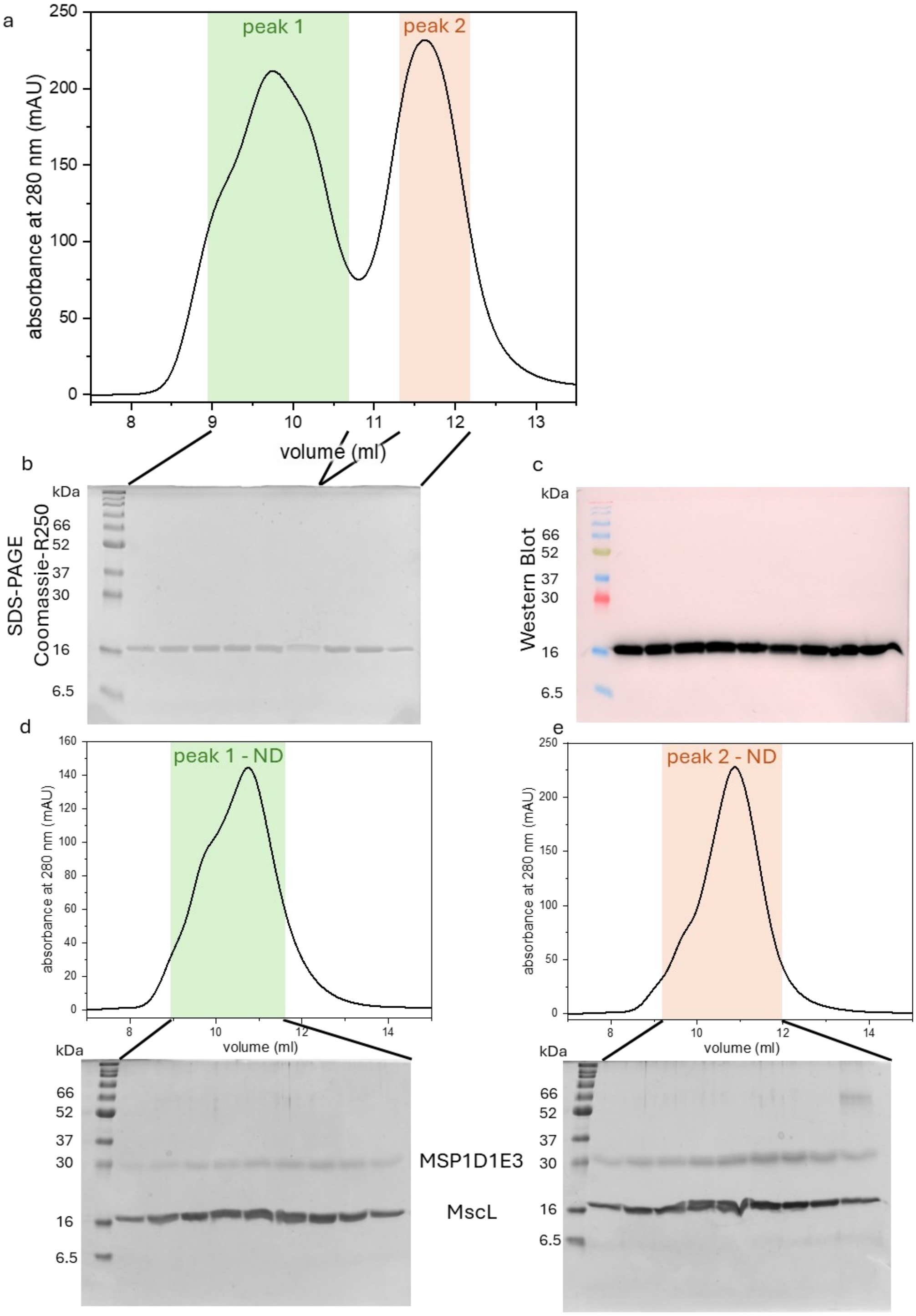


**Figure S1: Purification and reconstitution of EcMscL. a** Size exclusion chromatogram of EcMscL as shown in Figure 1 together with **b** SDS-PAGE of the peak fractions with Coomassie staining and **c** Western blot analysis against the His-tag of the same fractions as in b. **d** peak 1 and **e** peak 2 were reconstituted into nanodiscs. SEC profiles (top) and SDS-PAGE (bottom) of the reconstitution are shown, respectively.

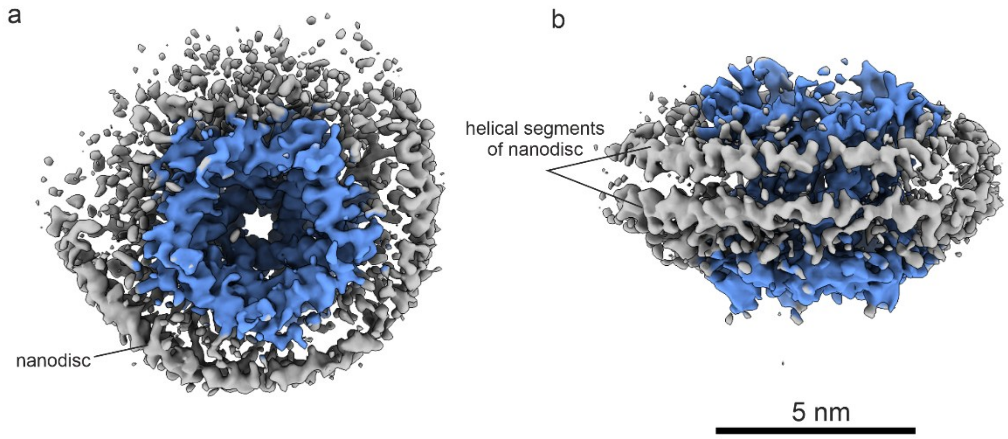


**Figure S2: Ab-initio reconstruction of EcMscL in nanodiscs**. (peak 1) with C1 symmetry. MscL is coloured in blue (color zone option of ChimeraX within 3.5 Å of the model) and the rest of the map in grey. Note, close to EcMscL, the helices of the scaffolding protein are resolved while further away from MscL the scaffolding is unresolved.

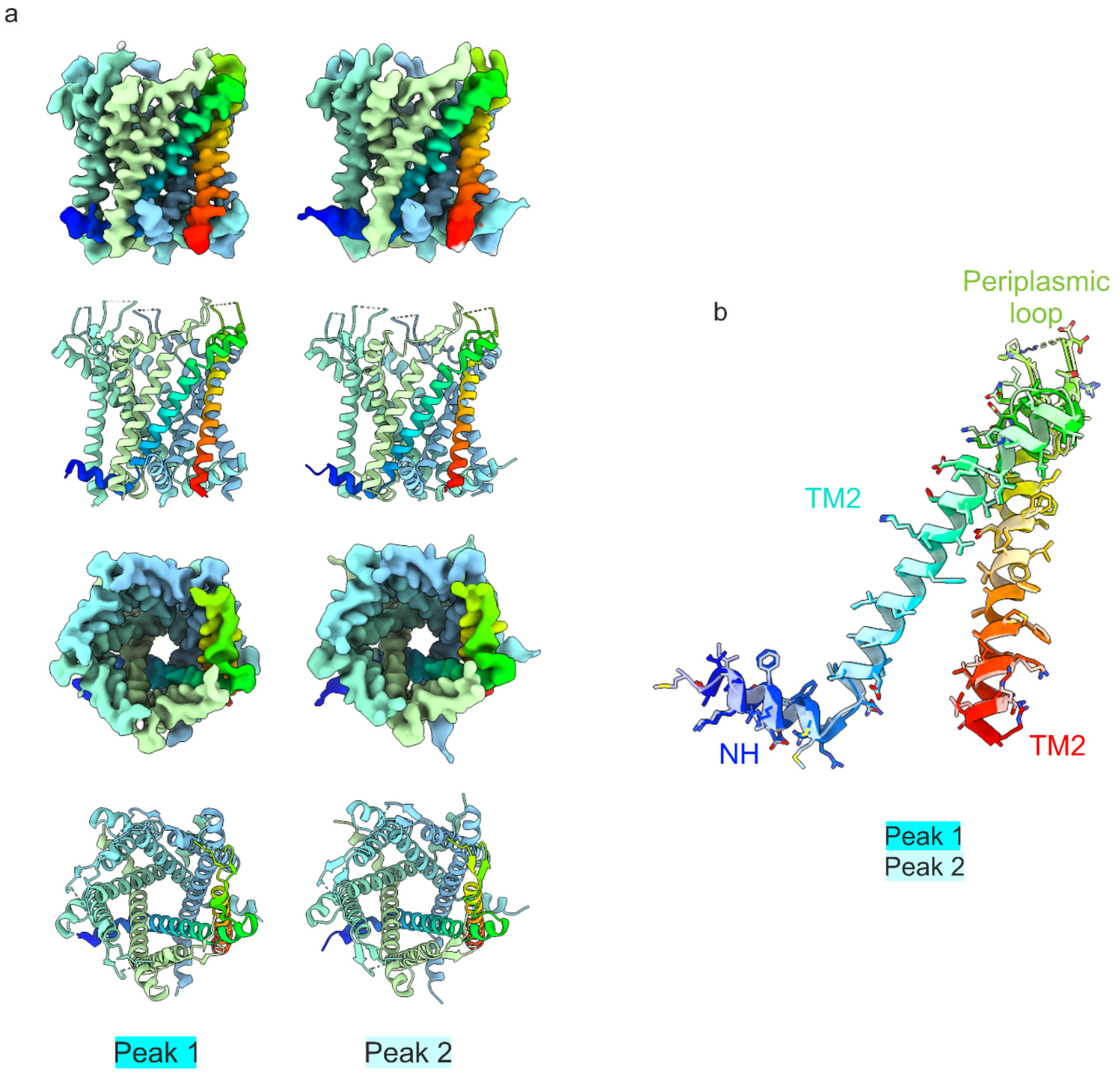


**Figure S3: Comparison of map and model of EcMscL of peak 1 and peak 2 reconstituted into nano-discs.** a) The maps (row 1 and 3) and models (rows 2 and 4) of EcMscL from peak 1 (left column) and peak 2 (right column) are shown. The surface representations of the maps are calculated at the same threshold. The surface of the maps is coloured with the same colours as in the models shown below (colour zone option of ChimeraX). Maps and models are presented in the same orientation. One chain is coloured in rainbow from blue (N-terminus) to red (resolved C-terminus). b) Cartoon representation of this chain in the same orientation as in the two top rows in a). The model derived from the reconstitution of peak 1 is shown in strong colours and of the reconstitution of peak 2 in pale colours. There is no obvious difference between the models.

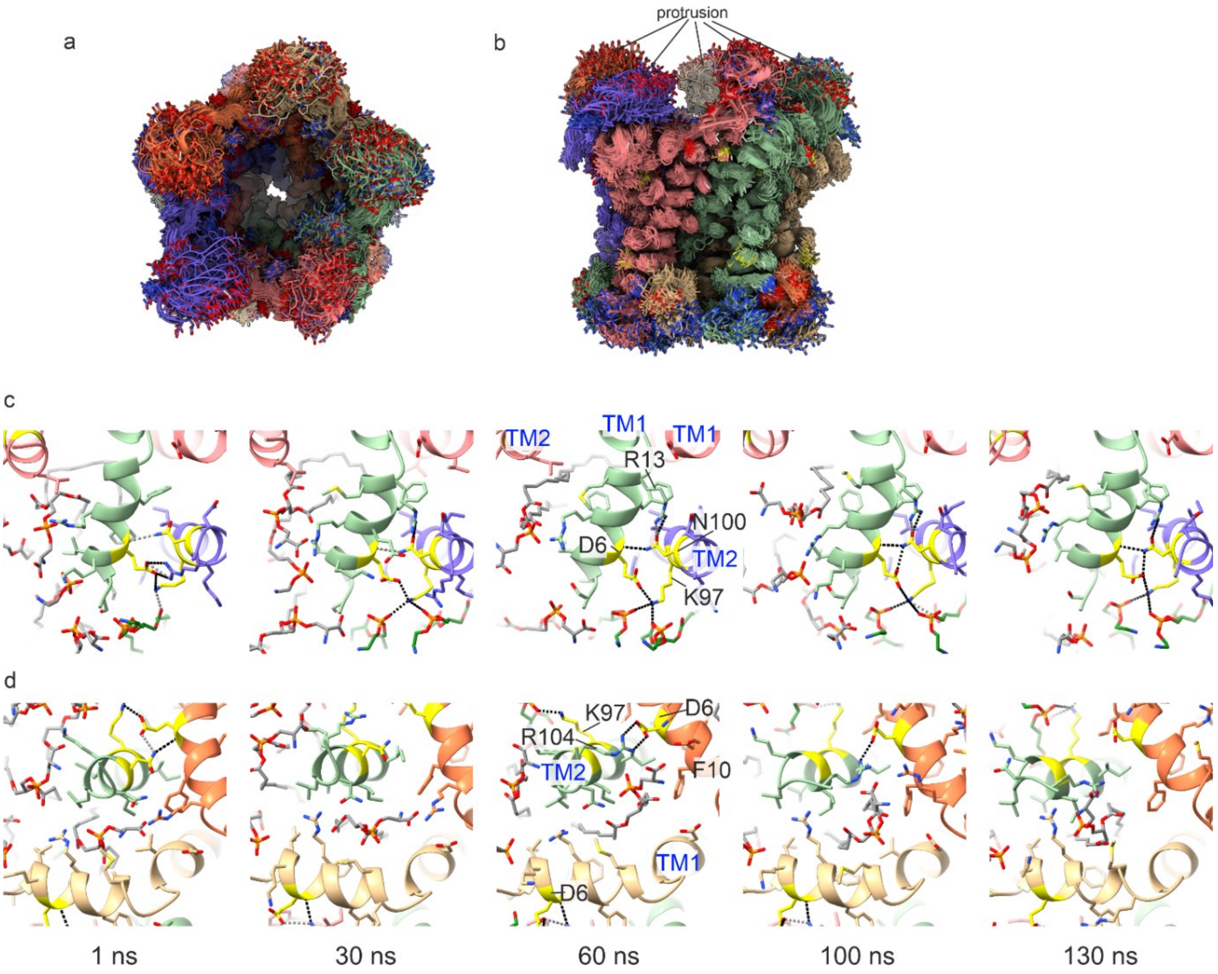


**Figure S4: Frames of the MD-trajectories**. a,b) Frames from a 200 ns trajectory shown every 2 ns. The frames are aligned by a least square fit of the protein. For clarity, water, lipids and KCl are omitted in the presentation. The protein included residues 2-104. The unmodelled residues 64-68 of the periplasmic loop were included in the simulation. Note, there is little divergence of the protein backbone and the side chains in the membrane part and there is large variability in the backbone of the periplasmic loop at the protrusions.

c,d) Close-up of a different trajectory at time points 1 ns, 30 ns, 60 ns, 100 ns and 130 ns. c) shows the interface between the N-terminal helix and the C-terminal helix involving residues E6, K97 and N100 (yellow). H-bonds between these three residues with other residues or lipids are shown as dotted, black lines. Lipids that were coordinated via an H-bond are shown in green. d) Interface on the other side of the N-terminal helix showing a lipid deeply penetrating the cleft. Lipids in contact with the N-terminal helix are shown in grey. Views in c) and d) are perpendicular to the plane of membrane and face the cytosolic side. Key residues are labelled in c) and d) in the respective 60 ns frames.

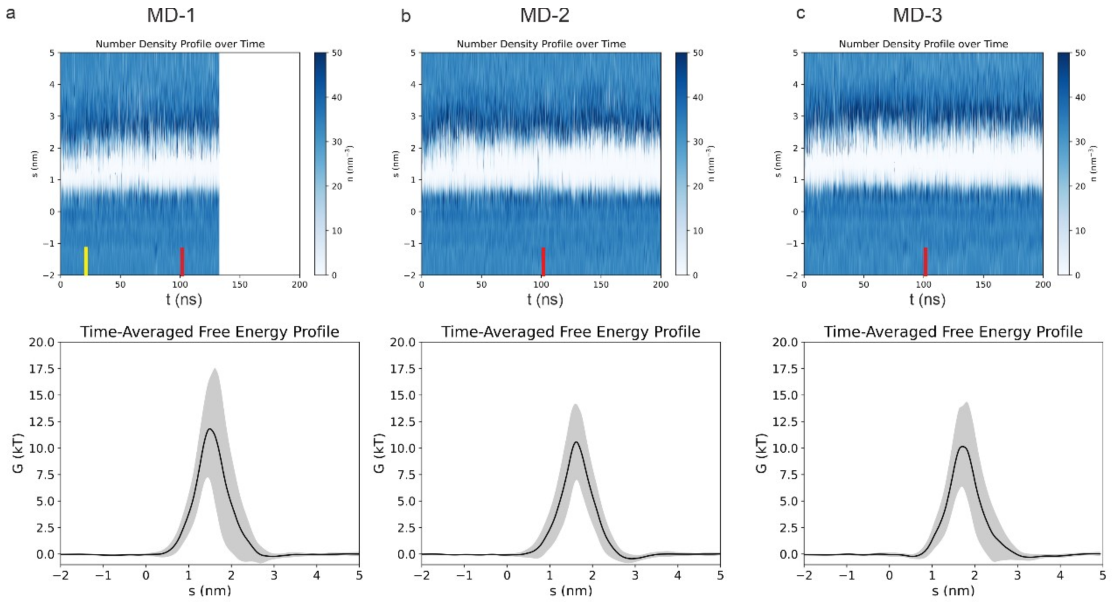


**Figure S5: Water density at the gate**. a-c) Three repeats of the MD simulation were calculated. The total simulation times were 130 ns (a) and 200 ns (b,c) The upper panels show the water density in the penetration pathway over the whole trajectory. The colour key corresponds to white for no water and dark blue for 50 water molecules/$nm^3$. For comparison: bulk water has a density of approximately 33.5 water molecules /$nm^3$. The lower panel shows the energetic barrier at the gate between 101 ns and 103 ns. This time interval is marked by red squares in the representation of the water density above. The yellow square in **a)** marks the time interval of 21 ns-23 ns for which the free energy was calculated that is shown in Figure 3c.

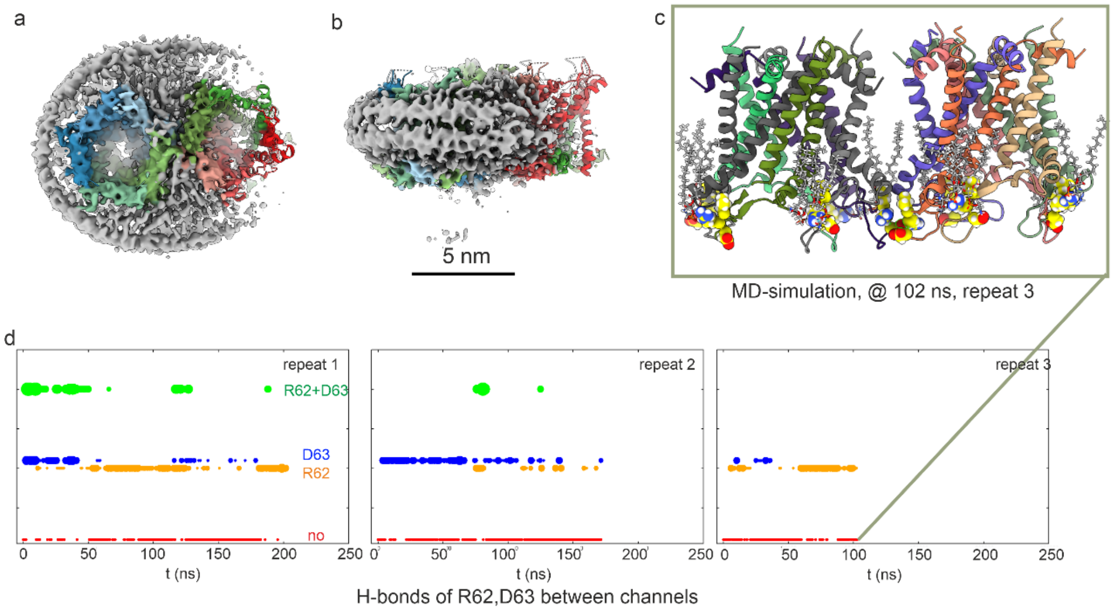


**Figure S6 Ab-initio map of dimer with C1-symmetry.** Two copies of the EcMscL model were placed as rigid body. The surface of the map was coloured according to the model within a zone of 3.5 Å. a) view from the periplasmic side, and b) rotated by 90°.

The rigid body dimer model was placed in a membrane and subjected to MD-simulation. **c)** The two monomers of the dimer stayed together for the whole length of the simulations. The example in c) shows the frame at 102 ns of repeat 3. The residues R62, D63 are colored in yellow and lipids coordinated by an H-bond via these residues in grey. R62 formed H-bonds to the head-groups of phospholipids in most chain except the ones at the contact side between the two channels.

**d)** Three repeats of the MD traces of different length (201 ns, 171 ns and 102 ns) were tested for H-bonds of R62, D63 of one channel with the same residues in the opposing channel. Red symbols indicate no H-bonds between the two channels, orange H-bonds between R62 and the opposing channel, blue between D63 and the opposing channel and green between R62 and D63 with the opposing channels. The size of the symbols correlates to the total number of detected H-bonds. H-bonds were detected with gromacs (option hbond)

All three repeats show that H-bonds formed quickly between both channels and fluctuated between the different possible bonding patterns indicating a dynamic but stable interaction between the channels.

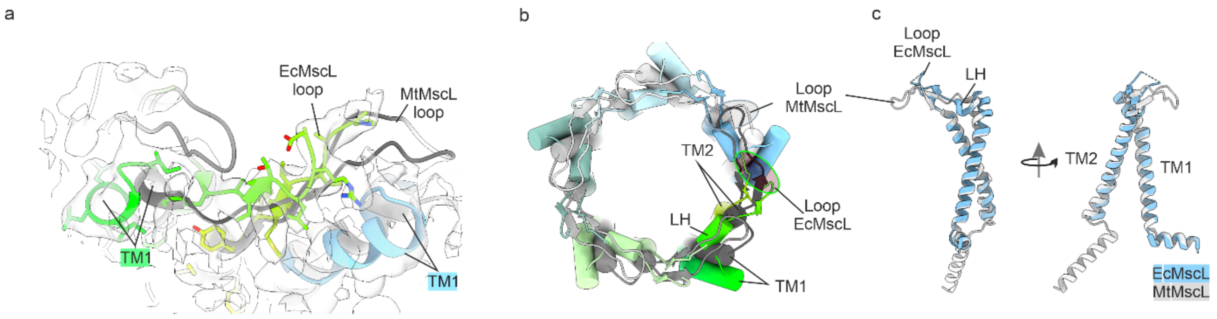


d

```
NH    TM1    LH    TM2

E.coli          MSIIKEFREFAMRGNVVDLAVGVIIGAAFGKIVSSLVADIIMPPLGLLIGGIDFKQFAVTLRDAQG
Mycobacterium   --MLKGFKEFLARGNIVDLAVAVVIGTAFTALVTKFTDSIITPLINR-IG-VNAQSDVGILRIGIG

cons              ::* *:**  ***:*****.*:**:**  :*:.:. .** * :.  ** :: :. .  **  . *

E.coli          DIPAVVMHYGVFIQNVFDFLIVAFAIFMAIKLINKLNRKKEEPAAAPAPTKEEVLLTEIRDLLKEQ
Mycobacterium   G--GQTIDLNVLLSAAINFFLIAFAVYFLVVLPYNTLRKKGEVE-QPGD-TQVVLLTEIRDLLAQT

cons            .  . .:. .*::. .::*:::***::: : *  :  *** *    *.  .: ********** :

E.coli          NNRS-----------------------
Mycobacterium   NGDSPGRHGGRGTPSPTDGPRASTESQ

cons            *. *
```

**Figure S7 Comparison of the model of EcMscL with the crystal structure of MtMscL** (pdb: 2OAR [3, 4]: **a)** Close up of the map at the periplasmic side with the fitted models of EcMscL (coloured, this study) and the crystal structure of MtMscL (grey, 2OAR [3, 4]). The models diverge in the orientation of the tips of the periplasmic loops and at the C-terminal end of TM1. **b)** View from the periplasmic side onto the cartoon representations of EcMscL (coloured, this study) and MtMscL (grey, 2OAR). The diverging ends of TM1, the position of the loop helix LH and the N-terminal end of TM2 are labelled. Note, the angled cross-section of EcMscL and the circular cross section of MtMscL on the periplasmic side. **c)** superposition of one subunit of MtMscL (grey) and EcMscL (blue). The two views are related by a 90° rotation around the vertical axis. Deviations map to the periplasmic loop and the C-termini of TM1 and TM2. **d)** Alignment of EcMscL and MtMscL. The helical segments in the EcMscL model are marked with coloured bars. The residues at the interface between two EcMscL in a dimer are marked by red triangles. The alignment was done with T-coffee [5]. Pink indicates good, yellow average and green bad conservation.

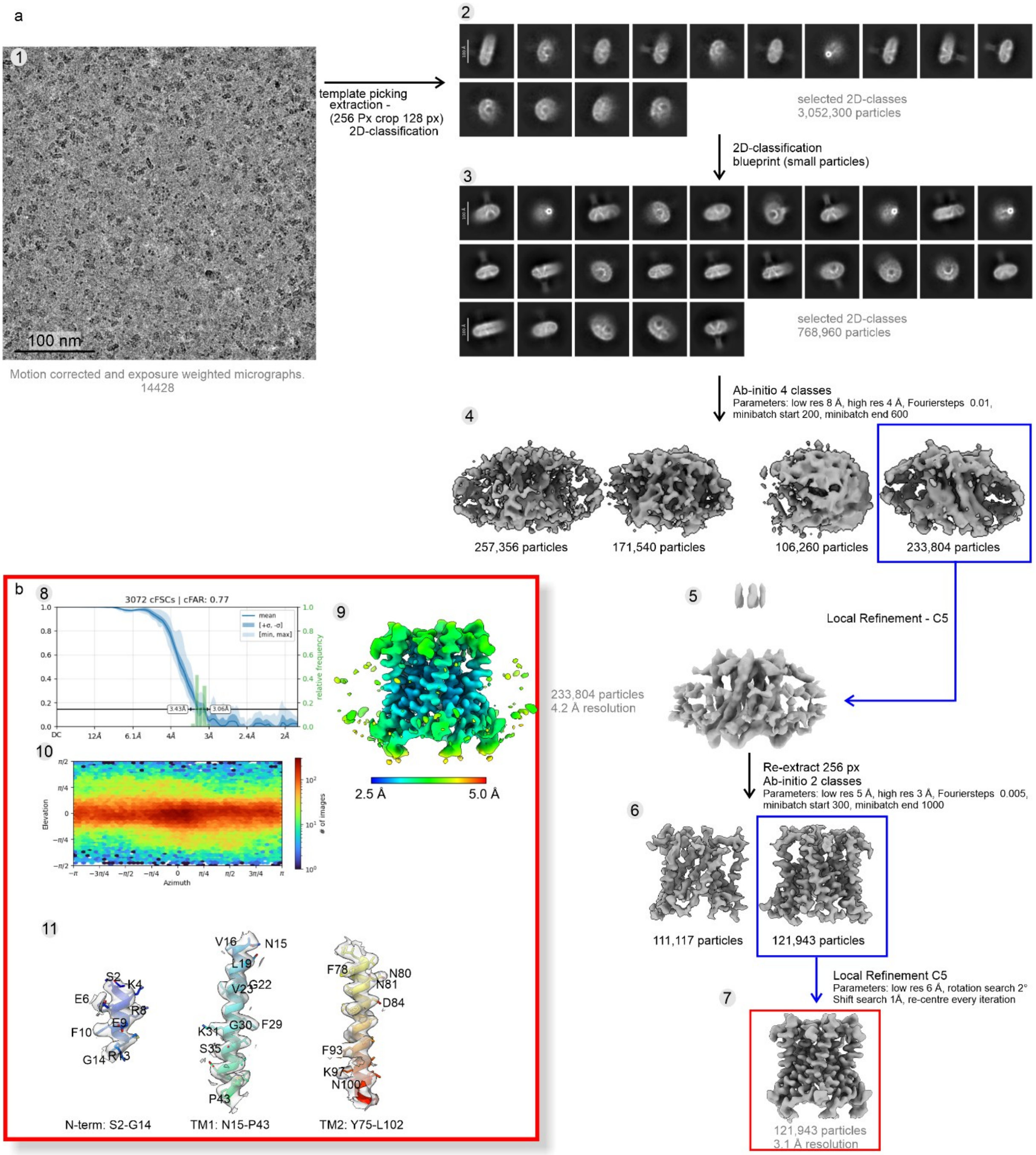


**Figure S8 Workflow of Image processing of EcMscL in nanodiscs (reconstitution of SEC-Peak 1)**. a) Workflow: Movies were motion corrected and exposure weighted during the live session in CryoSPARC [6]. Downstream imaging processing used the exposure-weighted movie averages (1). Particles were selected with a template derived from a 3D-volume of an ab-initio model. This model was generated during initial processing of blob-picked particles, was aligned to the C5 symmetry axis and reconstructed with C5 symmetry. The template selected particles were extracted in a 256x256 px² box Fourier-cropped to 128x128 px². The extracted particles were 2D-classified. The particles represented by the best class averages (2) were selected for a 2nd round of 2D-classification using the small particle blueprint. Again, the particles represented by the best class averages (3) were retained for further processing. The selected particles were processed by an ab-initio reconstruction with C1-symmetry into 4 classes (4). The best class (outlined by a blue square) comprised 233,804 particles and resolved the TM helices. Particles from this class were aligned to the C5 symmetry axis and mirrored. The particles were further refined in a local refinement with imposed C5 symmetry (5). Then, the particles were reextracted in a 256x256 px² box correcting the origins for the accumulated shifts. The re-extracted particles were subjected to a high-resolution ab-initio refinement [7] into two classes without imposing symmetry (6). The best class (blue outline) was subjected to local refinement with imposed C5 symmetry. The final map (7) had a resolution of 3.1 Å.

b) Evaluation of the final map with CryoSPARC: The conical Fourier Shell correlation (8) showed a resolution range of 2.9 – 3.6 Å. The map coloured by the local resolution (9) showed that the map was best resolved in the transmembrane part. The colour key is given below. The length of the colour key corresponds to 5 nm. The sampling of orientations in the final map is shown in (10). The model of the N-terminal helix, TM1 and TM2 are shown together with the surrounding map (11) to highlight the quality of the map. Some key residues are indicated. Note, the map is FSC-filtered but not B-factor sharpened.

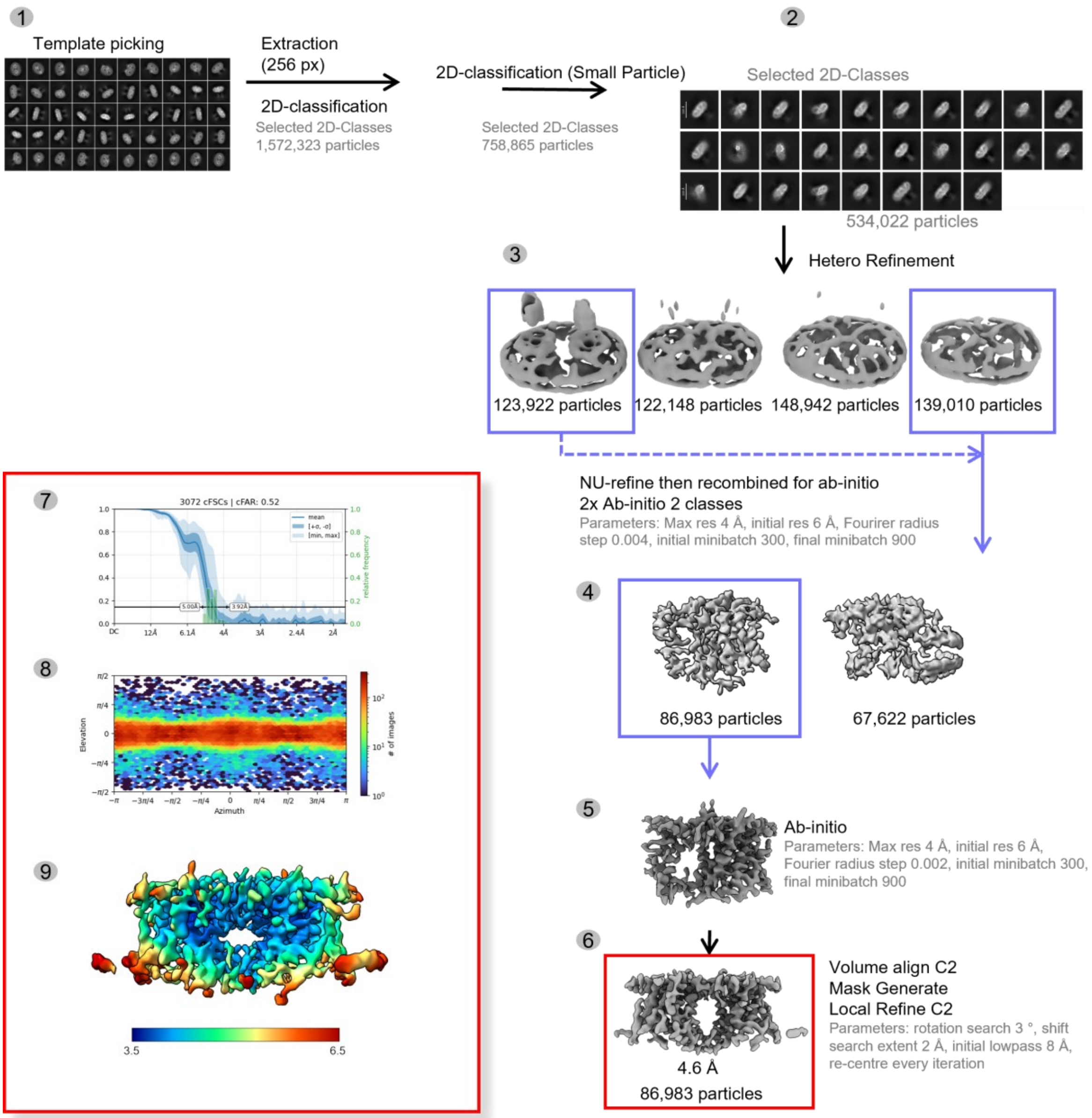


**Figure S9 of Image Processing of the EcMscL-Dimer**. 1)Template picking starts with the pre-processed movies (Figure S8) of EcMscL in nanodiscs from the reconstitution of SEC-Peak 1. (1) Templates were created from a 3D volume in cryoSPARC and 2,718,893 particles were extracted at a box size of 256x256 px². (4) After 2 rounds of 2D-Classification including one round using the small particle parameter blueprint on cryoSPARC 534,022 particles distributed across 28 classes remained. (3) The particles were used for heterogeneous refinement across 4 classes. The 4 classes were composed of 4 of the same volume that was used to generate the 2D-templates. This volume displaying a dimer conformation was generated as a class from a previous refinement strategy (Figure S8). (4) Two classes totalling 261,707 particles were first tested for NU-refinement before being recombined into one particle stack and used for ab-initio. The best classes (outlined by blue squares) were selected because of strong dimer features. (5) Ab-initio reconstruction was performed twice each time with two classes and the following parameters: enforce non-negativity off, initial resolution 6 Å, max resolution 4 Å, Fourier radius step 0.004, centre structures in real space off and larger initial and final minibatches. The best class after each ab-initio round was selected for strong helical features with the final class selected having 86,983 particles. (5) The particles from the best class were then used for a single class ab-initio reconstruction with similar parameters as (4) but with a Fourier radius step size of 0.002. (6) The resulting volume was then aligned to C2 symmetry and used to generate a mask. The particles were then further refined using local refinement with an initial lowpass of 8 Å, and C2 symmetry with a final resolution of 4.6 Å. (7-9) Evaluation of the final map with CryoSPARC. (7) The conical Fourier Shell correlation displayed a resolution range of 3.9 – 5.0 Å. (8) The sampling of orientations in the final 4.6 Å map. (9) The central helices between the dimers were best resolved as displayed by the local resolution. One dimer (left) showed worse resolution in the periphery helices. This was observed throughout refinement. The colour key is given below.

**Table S1 Cryo-EM data collection, refinement and validation statistics**

| | Pentamer in ND (EMDB-56958) (PDB 28YC) | Dimers of pentamers im ND ( EMD-58096) |
|---|---|---|
| **Data collection and processing** | | |
| Magnification | 130,000 | |
| Voltage (kV) | 300 | |
| Electron exposure (e–/Å$^2$) | 70 | |
| Defocus range (μm) | 0.3-2 | |
| Pixel size (Å) | 0.946 | |
| Symmetry imposed | C5 | C2 |
| Initial particle images (no.) | 6,478,112 | 2,718,893 |
| Final particle images (no.) | 121,943 | 89,983 |
| Map resolution (Å) | 3.2 | 4.6 |
| 0.143 FSC threshold | | |
| Map resolution range (Å) | 2.9-3.6 | |
| In cFSCs | 3.1-3.4 | 3.9-5.0 |
| **Refinement** | | |
| Initial model used (PDB code) | -- | |
| Model resolution (Å) | 3.5 | |
| FSC threshold | 0.5 | |
| Model resolution range (Å) | -- | |
| Map sharpening *B* factor (Å$^2$) | -50 | |
| Model composition | | |
| Non-hydrogen atoms | 3785 | |
| Protein residues | 490 | |
| Ligands | 0 | |
| *B* factors (Å$^2$) | | |
| Protein | 123 | |
| Ligand | -- | |
| R.m.s. deviations | | |
| Bond lengths (Å) | 0.007 | |
| Bond angles (°) | 0.800 | |
| Validation | | |
| MolProbity score | 1.76 | |
| Clashscore | 8.69 | |
| Poor rotamers (%) | 0.00 | |
| Ramachandran plot | | |
| Favoured (%) | 95.74 | |
| Allowed (%) | 4.26 | |
| Disallowed (%) | 0.00 | |
